# A CAR RNA FISH assay to study functional and spatial heterogeneity of chimeric antigen receptor T cells in tissue

**DOI:** 10.1101/2020.08.21.260935

**Authors:** Karsten Eichholz, Alvason Zhenhua Li, Kurt Diem, Michael C. Jensen, Jia Zhu, Lawrence Corey

**Affiliations:** Vaccine and Infectious Disease Division, Fred Hutchinson Cancer Research Center, Seattle, WA, USA; Department of Laboratory Medicine, University of Washington, Seattle, WA, USA; Clinical Research Division, Fred Hutchinson Cancer Research Center (FHCRC), Seattle, WA, USA; Ben Towne Center for Childhood Cancer Research, Seattle Children’s Research Institute, Seattle, WA, USA; Department of Medicine, University of Washington, Seattle, WA, USA

**Keywords:** Chimeric antigen receptor T cells, quantitative microscopy, histo-cytometry

## Abstract

Chimeric antigen receptor (CAR) T cells are engineered cells used in cancer therapy and are studied to treat infectious diseases. Trafficking and persistence of CAR T cells is an important requirement for efficacy to target cancer. Here, we describe a CAR RNA FISH histo-cytometry platform combined with a random reaction seed image analysis algorithm to quantitate spatial distribution and in vivo functional activity of a CAR T cell population at a single cell resolution for preclinical models. In situ, CAR T cell exhibited a heterogenous effector gene expression and this was related to the distance from tumor cells, allowing a quantitative assessment of the potential in vivo effectiveness. The platform offers the potential to study immune functions of genetically engineered cells in situ with their target cells in tissues with high statistical power and thus, can serve as an important tool for preclinical assessment of CAR T cell effectiveness.

**Brief summary:** We developed an imaging platform and analysis pipeline to study large populations of engineered cells at a single cell level in situ.

**One Sentence Summary:** We developed a CAR RNA FISH assay to study chimeric antigen receptor T cell trafficking and function in mouse tissue.

## Introduction

Chimeric antigen receptors (CAR) T cells are engineered T cells that are under development to treat various cancers and infectious diseases. The CAR is a recombinant fusion protein that combines an extracellular antigen-specific single chain variable fragment (scFv) with a transmembrane domain and an intracellular signaling domain to promote targeted killing. The scFv is directed against extracellular antigens, provides high specificity and induces cytolysis when the CAR T cell interacts with its target. Recently, three CAR T cell therapies based on the anti-CD19 scFv clone FMC63 have been approved for treatment of B-cell acute lymphoblastic leukemia (BALL) and non-Hodgkin lymphoma and numerous constructs are in pre-clinical and clinical development for a variety of hematological and solid tumors.

While CAR T cell therapies have shown some success in hematological cancers, the application in solid tumor settings is still hampered by the immunosuppressive tumor microenvironment, poor tumor infiltration, the choice of an efficient scFv-tumor antigen pair and adverse effects like cytokine release syndrome *(1–3)*.

CAR T cell performance, proliferation and persistence in preclinical *in vivo* studies and clinical application are mainly studied by flow cytometry, quantitative PCR, and for preclinical cancer mouse models only, whole body bioluminescence imaging of the tumor burden. These techniques are favorable to measure CAR T cell phenotype, functionality and expansion with high statistical power and specificity and can assess global anti-tumor efficacy, however, the location of the CAR T cells in the tissue with respect to the tumor cells cannot be determined.

Recently, RNA *in situ* hybridization (RNA ISH) approaches have been successfully applied to track CAR T cells in patients with glioblastoma *(4)*, in a case study in a patient with B-ALL, who died 5 days post-anti-CD19 CAR T cell infusion*(5)* and in clinical samples from the ZUMA-1 trial that tested the anti-CD19 CAR Axicabtagene ciloleucel *(6)*.

Automated image analysis requires reliable image segmentation, however, commonly used commercial and open source software mainly rely on nuclei detection and watershed algorithms. The weak or discontinuous cell membrane staining, overlapping cells tissue- or fixation-derived autofluorescence and other fluorescent artifacts have limited the use of current software packages for automated image analysis. To overcome the limitations, stochastic algorithms that use hidden-Markov-model (HMM) statistics proved to be useful to tackle the more advanced requirements of automated image analysis in tissue *(7–11)*. More recently, we developed a robust and efficient Random-Reaction-Seed method (RRS) that employs the maximum likelihood method to estimate probabilistic functions of Markov chains which showed efficient tracking of neurite net-like structures in fluorescence microscopy images *(12)*.

This article describes a RRS-assisted RNA fluorescence *in situ* hybridization assay to assess the spatial distribution and phenotype tumor cells and anti-CD19 CAR T cells in tissues of preclinical animal models. We use this technique to detect CAR T cells localization in tissue in a quantitative manner in a preclinical animal model. Our data indicate that proximity to tumor tissue influences in vivo killing potential as defined by *in situ IFNγ* and *GZMB* mRNA expression; thus, providing direct *in situ* evidence of in vivo activity and illustrating the importance of spatial interactions between the tumor microenvironment and in vivo functional activity of adoptively transferred T cells. The CAR RNA FISH imaging platform can be easily modified for novel CAR-antigen pairs to study the interaction between CAR T cell and the infiltrated tumor microenvironment and provide insights in determinants of potential CAR T cell effectiveness in preclinical models of this emerging cellular therapy platform.

## Material and methods

### Cell culture

TM-LCL and Jurkat were cultured in RPMI1640 (Gibco), 10% fetal bovine serum and penicillin/streptomycin.

### CAR T cells expansion

Anti-CD19 CD8 CAR T cells were cultured with TM-LCL feeder cells as previously described *(13)*. Briefly, 10^6^ anti-CD19 CD8 CAR T cells were expanded in presence of 7 x 10^6^ irradiated (8000 cGy) TM-LCL feeder cells in RPMI1640 containing 50 IU/mL human recombinant IL-2 (Peprotech), 10 ng/ml recombinant human lL-15 (Peprotech), 10% fetal bovine serum, 50 μM β-mercaptoethanol (SigmaAldrich) and 25 mM Hepes, pH 7.4 (Gibco) and penicillin/streptomycin. The anti-CD19 CD8 CAR T cells were used only for the *in vitro* probe validation.

### Tissue samples

The mouse tissues used for the CAR RNA FISH in situ validation contain a CD19+-BE2 cell neuroblastoma xenograft infused with anti-CD19 CAR T cells or an uninfused control tumor and were described in a previous study *(14)*. Briefly, the mouse received 50 million CD19 CAR T cells at a 1:1 CD4:CD8 ratio (25 million of each), and the tumor was harvested at day 7 post T cell infusion. The tumor was allowed to engraft for 10 days before T cell infusion in mice that did or did not receive CAR T cells later. The samples were provided by Michael C. Jensen and the actual in vivo experiment was not part of this study. They were chosen because they contain a mixture of human anti-CD19 CD4 and CD8 CAR T cells. The mouse tumor model was conducted under the Seattle Children’s Research Institute Institutional Animal Care and Use Committee (IACUC)-approved protocol.

### Flow cytometry

Day 12 CD8 CAR T cells TM-LCL co-cultures were stained with LIVE/DEAD Fixable Aqua Dead Cell Stain (ThermoFisher) followed by antibody staining against CD8 (FITC, clone RPA-T8, BD, 555366), CD19 (BV786, clone SJ25C1, biolegend) and EGFRt (PE, clone erbitux, provided by Juno Therapeutics) and analyzed on a LSR II flow cytometer. Gating on CD8+ EGFRt+ cells was done with fluorescence minus one-controls for CD8 and EGFRt.

### CAR RNA FISH - probes

zz probes for CAR mRNA were custom-made by Acdbio and are directed against the scFv, the area between the 41bb and CD3 signaling domain and in some cases the 3′UTR for CAR detection in tissues that are devoid of any other lentiviral constructs that may harbor the wPRE (Fig. 1A, Fig. 1 suppl.). The individual probes were compared *in silico* with the human, mouse and macaque (*M. mulatta* and *M. nemestrina)* genome to avoid cross-detection of endogenous counterparts of the CAR (e.g. CD3zeta).

**Figure 1:**
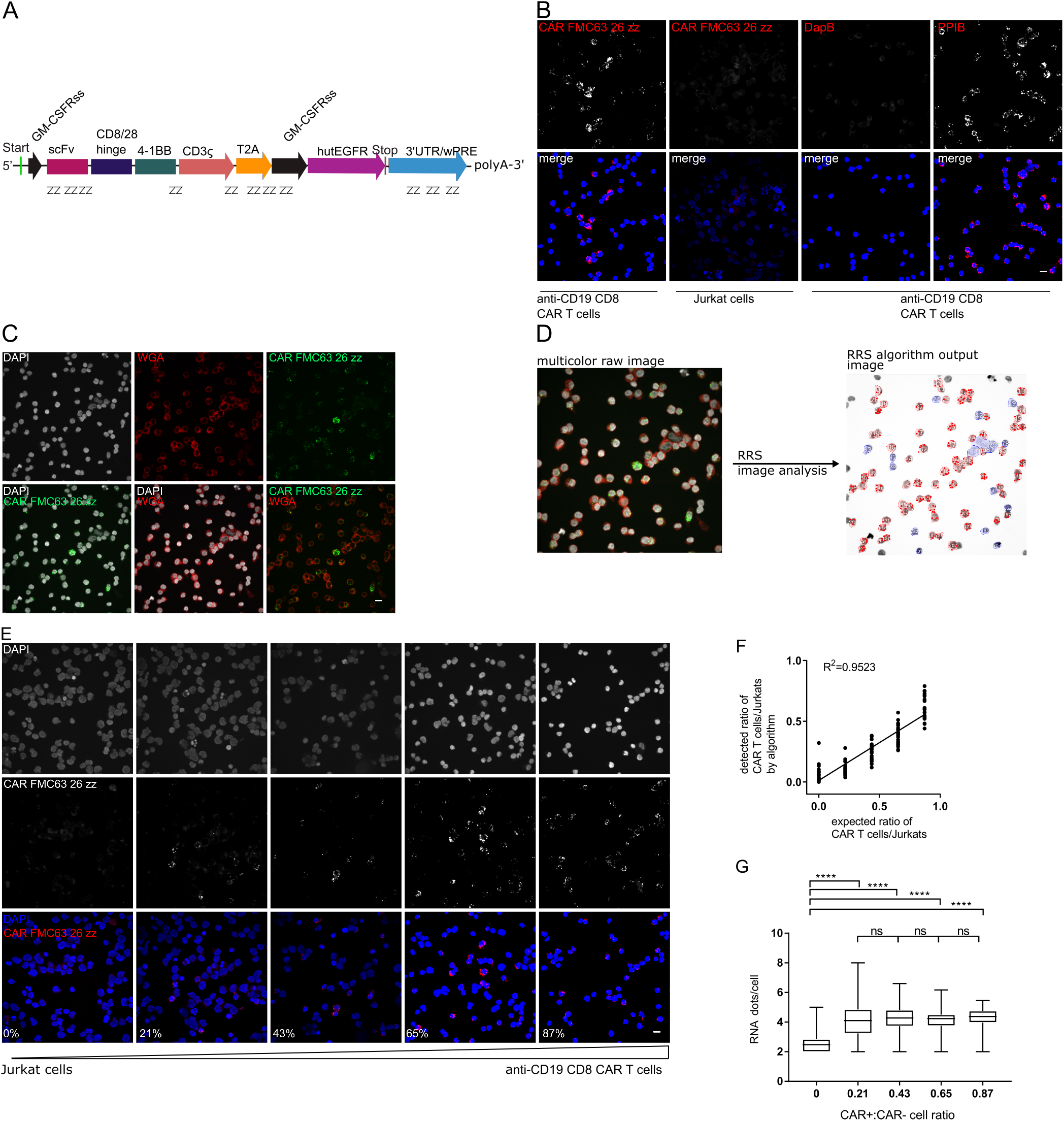
CD19 CAR RNA FISH *in vitro* validation. The probes for CAR mRNA detection are directed against the scFv and wPRE and bridging areas in the signaling domain of the CAR (A). Cytospins of actively growing human anti-CD19 CD8 CAR T cells or Jurkat cells were stained with CAR FMC63 26zz probes or PPIB (+ control) or DapB (- control) probes (B). CD19 CAR T cells were stained with CAR FMC63 26zz probes-Alexa488 and Texas red-conjugated wheat germ agglutinin to improve cell segmentation (C). RRS analysis of multicolor raw images includes nuclei and RNA detection as well RRS detection of plasma membrane staining (D). Fixed human anti-CD19 CD8 CAR T cells and Jurkat cells were mixed at different ratio, processed for cytospins and stained with CAR FMC63 26zz probes-Atto550, Oregon Green 488-conjugated WGA (not shown) and DAPI (E). Correlation of CAR+ cells detected by RRS analysis and flow cytometry (F). Comparison of RNA dots/cells between different dilution steps. Statistics are calculated by one-way ANOVA with Tukey’s multiple comparison test (G). Staining experiments were performed at least 3 times with technical duplicates. Scale bar = 10 μm

### CAR RNA FISH sample preparation - cytospins

To prepare cytospin slides to validate the probe specificity, 10^7^ anti-CD19 CD8 CAR T cells or Jurkat cells were washed twice with 40 mL PBS and centrifuged at 250 x g for 10 min. The cell pellets were resuspended in 10 mL 4% paraformaldehyde, PBS, fixed at 37°C for 1 h, and washed two times with 10 mL PBS. The cell pellet was then resuspended at 10^6^ cell/mL in 70% ethanol and stored for up to 7 days for further cytospin processing.

To obtain different densities of CAR T cells, CAR T cells were mixed with Jurkat cells at different ratio (0 – 100%). The cell suspensions were then centrifuged onto RNAse-free Superfrost plus glass slides (Electron microscopy sciences) with a cytospin centrifuge (cytospin 3, ThermoShandon) set to 800 rpm for 10 min to prepare the cytospins.

The glass slides were air dried for 20 min at ambient temperature and dehydrated stepwise in 50%, 70% and absolute ethanol for 5 min and stored in absolute ethanol at −20°C until further processing.

### CAR RNA FISH sample preparation - tissues

Fresh frozen optimal cutting temperature (OCT) compound-embedded tissues blocks were sectioned to 10 μm thickness onto RNAse-free Superfrost plus glass slides (Electron microscopy sciences) on a Leica CM1950 cryostat at −20°C. The slides were kept at −20°C for 1 h and transferred to −80°C or directly processed for FISH.

OCT sections were incubated in 4% PFA, PBS pre-chilled to 4°C for 15 min, washed 2 times briefly with PBS and dehydrated in a graded series of ethanol for 5 min each (50%, 70%, 2 x 100%).

Formalin-fixed, paraffin-embedded (FFPE) tissue blocks were sectioned to 4 μm thickness on a Leica RM2255 microtome. Sections were heated at 60°C for 1 h, dewaxed and processed for RNAscope pretreatment on a Leica Bond RX according to manufacturer’s recommendations with Leica Bond Epitope Retrieval Solution 2 (ACDbio) for 15 min at 95°C. Subsequently, FFPE sections were processed manually as described in the following section.

### CAR RNA FISH procedure

Cytospin slides or tissue section slides were removed from 100% ethanol after dehydration and air-dried for 30 min at 40°C in a hybridization oven or 5 min at ambient temperature, respectively. A hydrophobic barrier was created around the sample on the slides, air dried for 1 min, treated with H2O2 for 10 min (for RNAscope^®^ Multiplex Fluorescent Assay version 2) followed with protease (30 min protease III at 40°C for cytospin samples, 15 min protease IV at ambient temperature for mouse tissue sections in OCT-embedded tissues, or 30 min protease plus at 40°C for FFPE tissues) (ACDbio).

After protease digestion, samples were washed twice in PBS and anti-RNA zz probes against CAR, *CD4, CD8, IFNγ, GZMB, PPIB* and *DapB* were hybridized at 40°C for 2 h (Table 1 suppl.). As a negative control, the slides were treated with 4 μg/mL RNAase A and 100 U/mL RNAse T1 (RNase A/T1 mix, ThermoFisher) for 30 min between the protease digestion and the probe hybridization step. For the signal amplification after the probe hybridization, we used initially the RNAscope Multiplex Fluorescent Assay version 1 with the fluorophores Alexa 488 and Atto 550 and in later experiments the RNAscope Multiplex Fluorescent Assay version 2 with the fluorophores Opal 570 and Opal 650 (PerkinElmer) according the manufacturer’s protocol for multiplex fluorescent in situ hybridization kit. The advantage of using the red and far red Opal dyes for the RNA detection is the lower autofluorescence in tissues in this part of the UV/Vis spectrum and general better signal to noise ratio. We added an additional counterstaining for the plasma membrane before the nuclear counterstaining or omitted the nuclear counterstaining completely. In brief, the sample slides were washed 2 times for 2 min with PBS, incubated with 10 μg/mL wheat germ agglutinin conjugated with Texas Red-X or Oregon Green 488 (Invitrogen) for 10 min at ambient temperature, washed again 2 times for 2 min with PBS before proceeding to the nuclear counterstaining step.

### Digital droplet PCR (ddPCR) detection of CAR T cells in tissues

gDNA was extracted with QiAmp DNA kit (Qiagen)and 10 μL of DNA were used for ddPCR. Primer sequences were as follows for *wPRE* (5′-CGCAACCCCCACTGGTT-3′ [wPRE forward], 5′-AAAGCGAAAGTCCCGGAAAG-3′ [wPRE reverse] and 5′-FAM-CCACCACCTGTCAGC-MGB-3′ [wPRE probe]) and the CAR ORF (5′-AGCTGCCGATTTCCAGAAGA-3′ [CAR forward], 5′-CCAGGTTCAGCTCGTTGTACAG-3′ [CAR reverse] and 5′-FAM-TCAGCAGAAGCGCCGACGCC-QSY-3′ [CAR probe])

The DNA ddPCR was used with ddPCR Supermix for probe (Biorad) and the PCR conditions were 95°C for 10 min, then 40 cycles at 94°C for 30 s, 60°C for 1 min followed by 98°C for 10min. The thermal conditions were 50°C for 60 min for reverse transcription, 95°C for 10 min, then 40 cycles at 94°C for 30 s, 60°C for 1 min for the ddPCR followed by 98°C for 10 min.

### Immunohistochemistry

Immunofluorescence staining was performed after the last RNA target was developed (step Amp4 in the RNAscope Multiplex Fluorescent Assay version 1, TSA amplification step in the RNAscope Multiplex Fluorescent Assay version 2), the tissue sections were washed with PBS and blocked with 5% donkey or goat serum (Jackson ImmunoResearch), 2% BSA (Gemini Bio-Products) +0.2% Triton X-100 (SigmaAldrich) in 5X Casein (Vector Laboratories) for 30 min, followed by incubation with the mouse anti-huCD19 antibody to stain tumor cells (dilution 1:100,Clone ID: UMAB103, origene) overnight at 4°C.

Slides were washed, incubated with secondary donkey anti-mouse IgG-Alexa 647 or Alexa 488 (1:100; Molecular Probes/ThermoFisher Scientific) or a goat anti-mouse BV480 (1:100; Becton Dickinson) for 1 hour at room temperature, and washed 2 times for 5 minutes in PBS + tween (0.05% v/v). Subsequently, samples were counterstained with DAPI and mounted with ProLong Gold antifade (Thermofisher).

### Image acquisition

Images were acquired with a PerkinElmer Ultraview Vox spinning disk confocal microscope equipped with 4 laser lines 405, 488, 561 and 640 nm and a Hamamatsu ImageEM EMCCD camera (512 x 512 pixels, 16 micron pixel size, 90% Quantum efficiency) with a 40 x objective, a Leica SP8 confocal microscope equipped with laser lines at 405 nm, 440 nm, and adjustable white light laser, or a Nikon eclipse Ti epifluorescence microscope equipped with a sola light machine, a Hamamatsu ORCA-Flash4.0 LT+ C11440 (2048 ×2048 pixels, 82 % Quantum efficiency) and Plan Apo 10x 0.45, Plan Apo 20x 0.75 and Plan Fluo 40x 1.3 objectives. Images that were analyzed with dlRSS algorithm were not modified post-acquisition unless otherwise stated in the description of the algorithm. Images for publication figures that were acquired with the spinning disk confocal microscope were size-adjusted to 2048 x 2048 pixels and z-stacks were projected with the Max intensity function in the Fiji software package *(15)*.

### CAR T cell detection algorithm

Imaging data was analyzed on a GPU workstation with a 32 core CPU, 64 GB RAM and a QUADPRO P5000 GPU. To perform image segmentation based on WGA-plasma membrane staining, we extended a previously described algorithm for identification of neurite net-like structures and to generate the ground truth automatically for each image *(12)*. It is a robust and efficient RRS algorithm that employs the maximum likelihood method to estimate probabilistic functions of Markov chain. A U-net-based auto encoder decoder convolutional deep neuronal network *(16)* performs image segmentation and phenotyping based on the presence of nuclei, RNA and/or protein.

To detect the cell membrane two subsequent steps are performed by the algorithm. First, it extracts a net-like draft of the cellular border based on the RRS results and second, it performs cell-loop segmentation of the net-like draft and thus, also removes false positive membranes from the draft. Each tile of a tile scan is analyzed separately and to avoid any redundancy of areas that overlap on the border of two adjacent images a 48-pixel wide area was omitted from the analysis on each side of the tile. This equals to (4 x 48 x 2048 pixels)/(2048 x 2048 pixels) = 0.09 or 9% of each slide that were not used for analysis. To overcome the optical vignetting effect in the corners of the images were normalized using a Gaussian function.

For RNA FISH and CD19 IHC detection, a conventional local-maxima method is used and are then combined with the membrane map.

For the comparison of *GZMB* and IFNγ expression in CAR T cells in the CD19high and CD19low areas, CD19high and CD19low areas were defined by generating a circle for each CD19+ cells that corresponds to the actual area covered by the cell at its x y position. The circles were expanded approximately 16 times the radius of the circle and connective groups were created by overlapping the expanded circles. The largest connective group was denoted as CD19high area whereas the remainder of the tissue section was defined as CD19low area. The spatial density of CAR+ *GZMB-*, CAR+ *GZMB+,* CAR+ IFNγ-, CAR+ IFNγ+ was than calculated for the CD19high and CD19low area of each individual tissue section.

The RNA and protein expression levels as well as the location of the cells was exported as a comma-separated values file and loaded into the flow cytometry data analysis software FlowJo 10 (Becton Dickinson). In FlowJo, the imaging data was analyzed with the gating and plotting tools. To compare cell counts of the dnnRRA and manual operators, manual operators used the cell counter function in imageJ to track cell counts.

### Statistical analysis

The data were analyzed with graphpad prism software with the statistical tests stated in the figure legends. One-way ANOVA combined with a Tukey test was used to compare multiple groups. *, **, ***, **** correspond to p < 0.05, p < 0.01, p < 0.001 and p < 0.0001, respectively. Paired t test was used to compare the spatial density within the CD19high and CD19low areas. *, **, ***, **** correspond to p < 0.05, p < 0.01, p < 0.001 and p < 0.0001, respectively.

Box and whiskers plots show the median, the 5th to 95th percentile for the whiskers and the 25th to 75th percentile for the box.

## Results

### CAR RNA FISH detects CAR T cells in *in vitro* cultures

We designed the CAR RNA probes to be directed against the wPRE in the 3′UTR and the scFv of the anti-CD19 CAR FMC63 *(17)* (Fig. 1A, Fig. 1 suppl.) and combined it with a recently published image segmentation and analysis algorithm *(12)*.

For *in vitro* experiments, T cells were co-cultured with γ-irradiated CD19-expressing TM-LCL lymphoblastoid cells for 12 days and purity was assessed by flow cytometry (~90%) (Fig. 2 suppl.). To validate the CAR FMC63 26ZZ probes, anti-CD19 CD8 CAR T cells or Jurkat cells were processed for cytospins and stained for CAR mRNA (Fig. 1B). The anti-CD19 CD8 CAR T cells stained positive with the CAR FMC63 26ZZ probe set. By contrast, Jurkat cells stained with the CAR FMC63 26ZZ probe set were negative for CAR RNA spots. RNA FISH against housekeeping gene *PPIB* (Peptidyl-prolyl cis-trans isomerase B) and the bacterial gene *DapB* as positive and negative controls, respectively, for RNA integrity and probe specificity, yielded characteristic RNA FISH staining only in cells with relevant genes (Fig. 1B).

RNA FISH-based imaging techniques detect the target mRNA mostly in the cytosol that surrounds the nucleus as demonstrated with the CAR mRNA (Fig. 3 suppl.). The density of tissue impedes automated image segmentation to define the quantity of T cells expressing the majority of the extranuclear CAR mRNA and cannot be defined based on the nuclear DAPI staining. Traditional algorithms rely on nuclei detection and assume a slightly extended perimeter for the cytoplasm to detect the RNA. To define cellular boundaries and to facilitate computational image segmentation, we used fluorescently labeled wheat germ agglutinin as a second counterstain to label N-acetyl-D-glucosamine and sialic acids on cell surface glycoproteins (Fig. 1C). This allowed to analyze the images with a modified version of a RRS algorithm recently published by our group *(12)* (Fig. 1D). In brief, the acquired images are partitioned and, several 1000 seeds are distributed on the WGA plasma membrane stain images (Fig. 1D). The algorithm searches for the plasma membrane high fluorescence intensity signal, follows the signal using hidden Markov model statistics and tries to close a loop, which would represent a cell and thus, creates a membrane map *(12)*. Once the cell circumference has been determined, the algorithm uses classical Laplacian of Gaussian local maxima detection for nuclear, IHC and RNA staining within this perimeter (Fig. 1D). The output data contains the location and RNA expression levels of each individual cell of the input image (Fig. 1D).

To validate the CAR RNA FISH approach in combination with the RRS algorithm, we stained anti-CD19 CD8 CAR T and Jurkat cells mixed at different ratio (1, 0.75, 0.5, 0.25 and 0) with the CAR FMC63 26zz probe set and acquired images. The number of cells stained positive by CAR RNA FISH and the number of cells detected with RRS increased with incrementing number of input anti-CD19 CD8 CAR T cells (Fig. 1E). The input CAR T cells were 90% EGFRt+ by flow cytometry (Fig 1A suppl.), hence, we expected to detect ~ 90%, 67.5%, 45%, 22.5% and 0% CAR+ cells for the corresponding ratio of 1, 0.75, 0.5, 0.25 and 0. The RRS algorithm detected ~5 - 60% CAR+ cells in the CD19 CAR T cell data set. There was a strong correlation between detection of the algorithm and the expected values (R^2^=0.95) (Fig. 1F) and a significantly higher number of CAR RNA dots/cells compared to Jurkat cells alone (Fig. 1G). Combined, these data indicate that the CAR RNA FISH probe design and RRS image analysis is highly specific and quantitative for detection of CAR T cells in cell culture samples.

### CAR RNA FISH detect CAR T cells *in situ* in mouse xenografts

To validate CAR RNA FISH for *in situ* applications in preclinical animal models, we used a BE2 neuroblastoma xenograft model of anti-CD19 CAR T cell-infused mice or control mice without CAR T cell infusion *(14)*. The BE2 cell line was transduced with multiple lentiviruses to express CD19 and luciferase and may contain the common wPRE sequence. We performed digital droplet PCR with primers directed against the signaling domain and the wPRE in genomic DNA. We detected the wPRE portion in both CAR+ and CAR-BE2 tumor tissue whereas the CAR signaling domain was only present in CAR+ BE2 tumor tissues (Fig. 4 suppl.). Therefore, to exclude false-positive cell detection due to cross-hybridization artifacts during the *in situ* setup of the CAR RNA FISH on mouse xenograft tissues, we only used a reduced probe set of total 15 zz probes, coined FMC63-15ZZ, and omitted the wPRE probes (Fig. 1 suppl.).

We observed a characteristic RNA FISH pattern with the CAR RNA FISH in formalin-fixed paraffin-embedded CD19+ BE2 xenograft tissues from mice infused with anti-CD19 CAR T cells but not from uninfused control mice (Fig. 2A). To ensure that mRNA integrity was of sufficient quality in the mouse xenograft and the RNA FISH staining procedure gave only low unspecific staining, we used probes against the house keeping gene *PPIB* (Fig. 5A suppl.) and the bacterial gene *DapB* (Fig. 5B suppl.), which gave a ubiquitous staining pattern or did not result in any positive signal in the corresponding area on adjacent slides, respectively. To further confirm the specificity of the CAR RNA FISH signal, no FISH signal could be observed when we omitted the FMC63 15ZZ probe set in the probe hybridization step (Fig. 5C suppl.), or after pretreatment of CAR T cell-positive tissues with RNAse A/H (Fig. 5D suppl.). These data indicate that the CAR RNA FISH approach can be applied for CAR T cell detection in formalin fixation and paraffin embedded tissues. We performed similar experiments in CAR T cell-positive and -negative mouse xenograft tissue section that went through the process of fresh-frozen OCT embedding and obtained similar results (data not shown).

**Figure 2:**
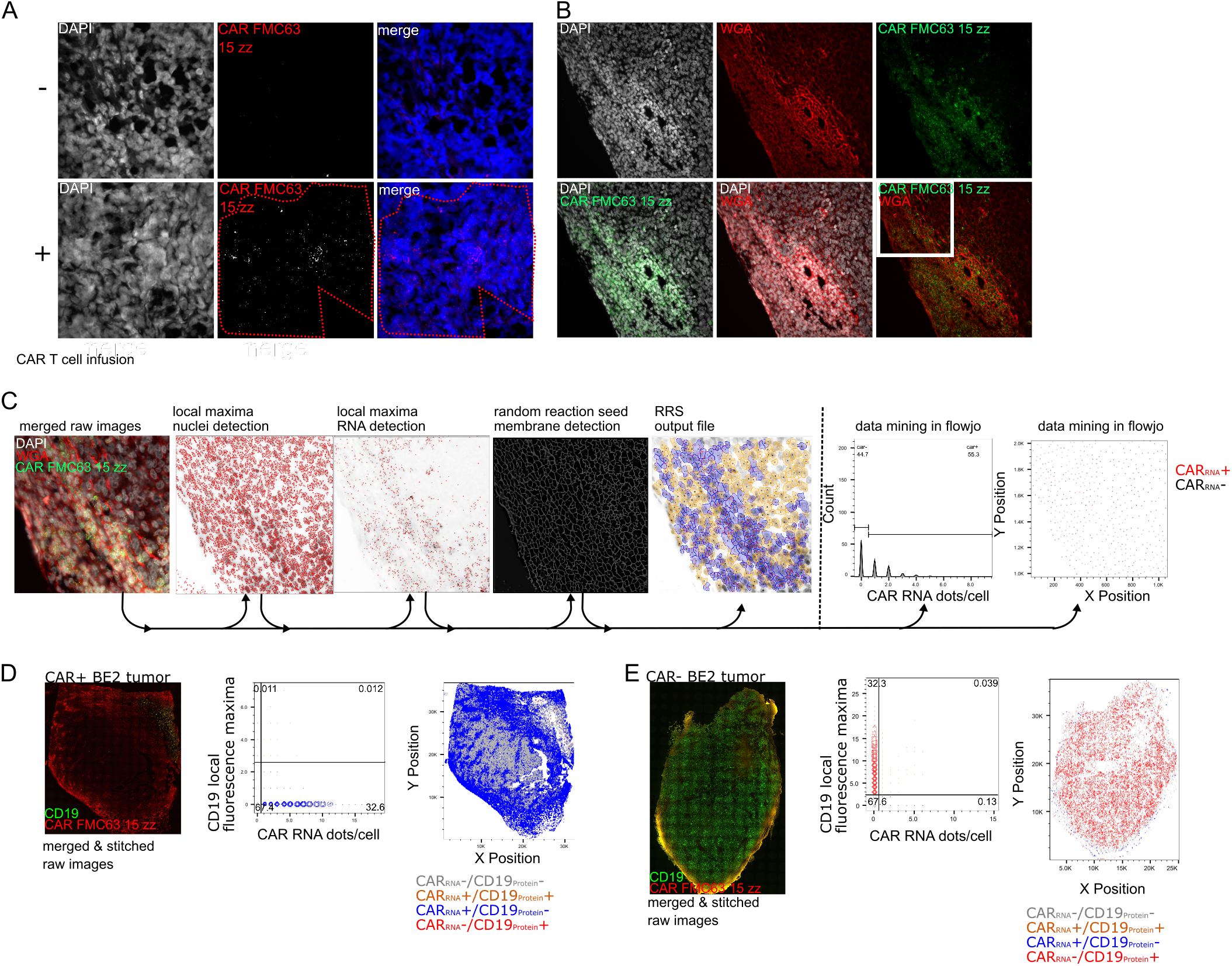
in situ detection of anti-CD19-CAR T cells in mouse xenografts of NSG mice after anti-CD19 CAR T cell infusion. FFPE-BE2 neuroblastoma xenografts with ectopic expression of truncated CD19 treated with anti-CD19 CAR T cell for 7 days or without CAR T cells was used for the CAR RNA FISH *in situ* validation. Staining of CAR+ and CAR-FFPE-BE2 tumor tissue sections with CAR FMC63 15zz probes (red). The red dotted line denotes an area that showed specific CAR RNA FISH staining (A). Staining of CAR+ BE2 tumor with CAR FMC63 15zz probes and Texas Red-conjugated wheat germ agglutinin improves cell segmentation (B). RRS analysis of multicolor raw images includes nuclei and CAR RNA detection as well RRS detection of plasma membrane staining. Data mining can be performed, and differential RNA expression and cell centers can be visualized with Flowjo. The example picture derives from the area framed in white in B (C). Whole tissue overview images show the distribution of the CAR RNA and CD19 IHC staining combined with WGA and DAPI (not shown) and the whole tissue RRS analysis of CAR+ (n=3) (D) and CAR-(n=3) (E) tumor section.

To quantify the RNA expression of the CAR and any other mRNA of interest on a per cell basis on the complete tissue section by RRS, we used WGA to reliably define the cell boundaries of each cell in the tumor section (Fig. 2B). As described before, the RRS algorithm uses local fluorescence maxima detection for nuclei and RNA and RRS WGA-based image segmentation to quantify the spatial distribution and frequency any mRNA of interest on a per cells and the data can be exported and analyzed in flow cytometry analysis software like Flowjo by the individual researcher (Fig. 2C). To exemplify the performance of the RRS algorithm for image segmentation and cell counting, we chose 10 images from our data set (Fig. 6A suppl.) that showed typical fluorescence intensities for nuclei and membrane staining for different imaging situations. The RRS algorithm and three operators counted the total cell number manually in imageJ based on membrane staining and nuclei staining (Fig. 6B suppl.). The observed values obtained by manual counting or RRS correlated across the example images (Pearson’s r<0.9 for each of the operators) (Fig. 6C suppl.). The average time required per image was 15:31 min ± 8:02 min for manual counting and 1:28 min ± 0:18 min for the RRS algorithm (Fig. 6D suppl.). In addition to the total number of cells, the RRS algorithm also segments and quantifies RNA FISH and IHC fluorescence data for T cell- or tumor-related genes, x-y position of the individual cell and cellular borders and integrates and stitches adjacent micrograph images. Notably, the image data sets recorded for the individual tissue sections frequently exceeded 300 multichannel images per sections and we were thus able to interrogate multiplexed mRNA expression levels in 150000 to 250000 cells per tissue section on a per cell basis.

We analyzed image data sets of a CAR RNA FISH RNA FISH combined with immunohistochemistry for CD19 from CAR T cell-positive and negative BE2 xenograft tissues with the RRS algorithm and used FlowJo for data analysis. We detected 32.6% and 0.13% CAR+ cells in the CAR+ (Fig. 2D) and CAR-BE2 (Fig. 2E) tumor, respectively. CD19 expression was below 1% and above 30% of cells in CAR+ BE2 tumor and CAR-BE2 xenograft (Fig. 2D, E), respectively.

These data indicated that the CAR RNA FISH detects specifically CAR T cells in tissues from preclinical *in vivo* models and that the RRS algorithm can be used to analyze CAR T cell abundance, spatial distribution and efficacy.

### CD4 and CD8 CAR T cells are widely distributed in the tumor and display differential functional and spatial gene expression

To further characterize the functionality of the CAR T cells, we then set out to do combinations of multiplex RNA FISH for the CAR, the T cell lineage marker genes *CD4* and *CD8α,* as well as the effector genes *IFNγ* and *GZMB* and analyzed the image data sets with the RRS algorithm to phenotype the tissue.

Independent CAR RNA FISH in combination with *CD4* or *CD8α* showed that the CD4 and CD8 CAR T cells are evenly distributed within the CAR+ foci (Fig. 3A, B). We then subjected the image data to computational analysis with the RRS algorithm. For the CAR and *CD4* RNA FISH, 17.2% of the total cells were CAR+/*CD4+* and 24.4% CAR+/*CD4-* (presumably CAR+/*CD8+* T cells) (Fig 3C). For the CAR and *CD8α* RNA FISH, 20.6% of the total cells were CAR+/*CD8α*+ and 19.9% CAR+/*CD8α* - (presumably CAR+/CD4+ T cells) (Fig 3D). Thus, in this model the tissue-based distribution was relatively similar between the CD4+ and CD8+ CAR T cells.

**Figure 3:**
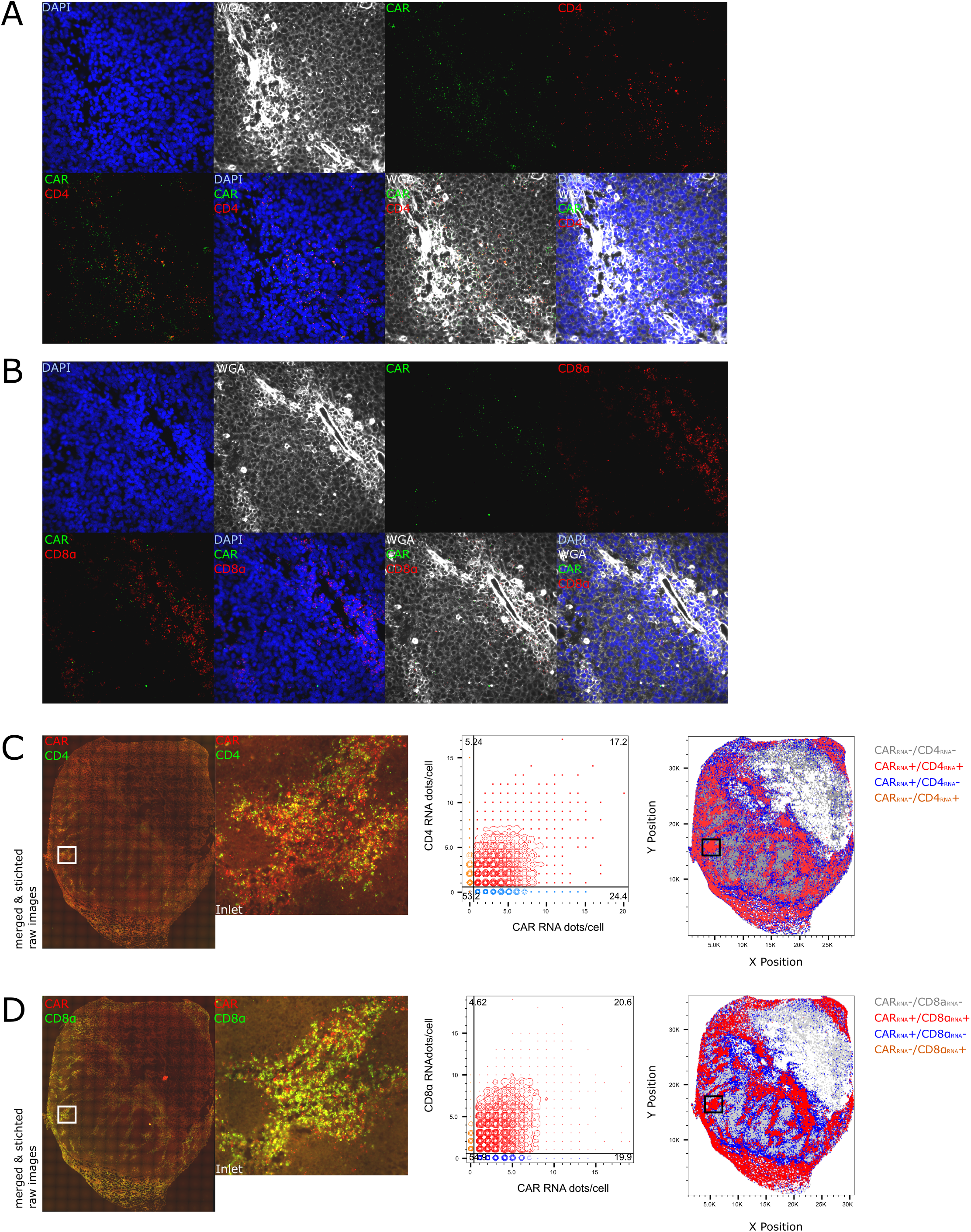
CD4 and CD8 CAR T cells are present in the tumor and express IFNγ. CAR T cell lineage phenotyping was done with CAR RNA FISH in combination with DAPI, WGA and *CD4* or *CD8α* RNA FISH on a CAR+ CD19+ FFPE BE2 neuroblastoma xenograft. Representative images from the tile scans are shown for CAR RNA and *CD4* RNA (A) or *CD8α* RNA (B) as well as nuclear and plasma membrane counterstaining. The single images from the tile scan were analyzed by RRS. The single tiles of the CAR (red) and *CD4* (green) RNA FISH channels were used to create whole tissue overview images to visually assess cells distribution and the whole tissue RRS analysis of the two RNA FISH, the WGA and DAPI channel of each single tile shows the distribution of the CAR+ CD4+ and CD4-T cells in a representative CAR+ BE2 tumor section (n=3) (C). The single tiles of the CAR (red) and *CD8a* (green) RNA FISH channels were used to create whole tissue overview images to visually assess cells distribution and the whole tissue RRS analysis of the two RNA FISH, the WGA and DAPI channel of each single tile shows the distribution of the CAR+ CD8+ and CD8-T cells in a representative CAR+ BE2 tumor sections (n=3) (D).

To further characterize the functionality and heterogeneity of the CAR T cell pool within the CAR+ BE2 tumor, we performed IHC for CD19, CAR RNA FISH combined with either *GZMB* or *IFNγ* RNA FISH on several sequential slides from the same tissue block. CAR+ cells that express high levels of *GZMB* RNA (Fig. 4A) or *IFNγ* RNA (Fig. 4B) can be observed by confocal microscopy close to the residual tumor. Concomitantly, CAR+ cells that are located in areas with low or absence of CD19 expression only express low levels of either *GZMB* or *IFNγ*RNA (Fig. 7 suppl. A and B). Whole-tissue tile scans of the CD19, CAR RNA FISH co-staining with either *GZMB* RNA FISH (n=5) (Fig. 4C) or *IFNγ* RNA FISH (n=5) (Fig. 4D) also indicated that the highest expression of *GZMB* but not *IFNγ* is in close proximity to the tumor. Examples images from this scan in close proximity of the residual tumor are also shown in the data supplement (Fig. 7 suppl. C and D). The data sets were analyzed by RRS and the x y position of CD19+ and CAR+ *GZMB*-/CAR+ *GZMB+* (Fig. 4E and Fig. 8 suppl.) or CAR+ *IFNγ*-/CAR+ *IFNγ+* (Fig. 4F and Fig. 9 suppl.) cell were plotted separately or combined in Flowjo.

**Figure 4:**
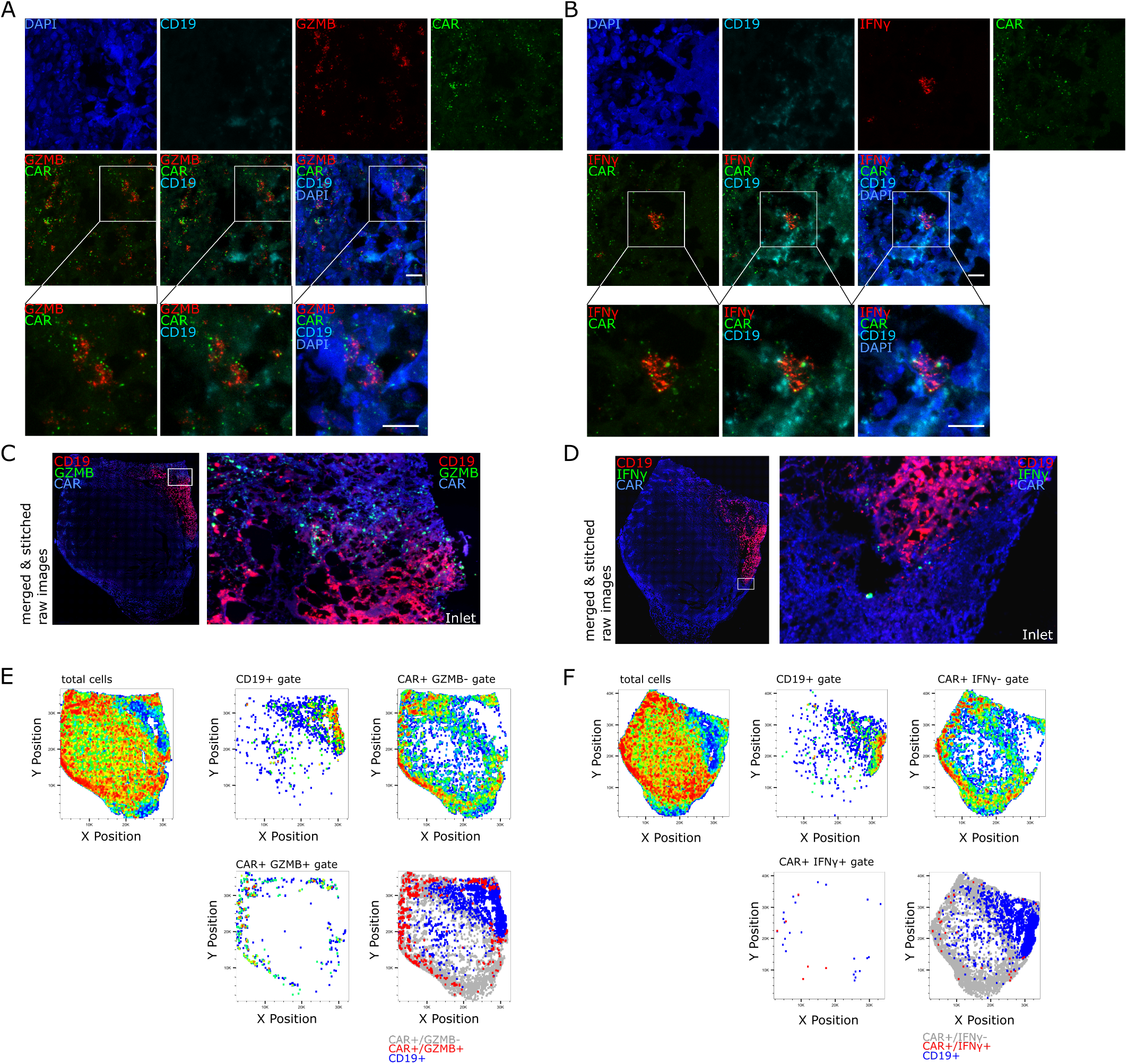
*GZMB* and *IFNγ* are highly expressed in a minority of CAR+ cells. CAR T cell functional phenotyping in a FFPE BE2 neuroblastoma xenograft was done by staining with DAPI, WGA, IHC for tumor marker CD19 and CAR RNA FISH in combination with either *GZMB* (n=5) or *IFNγ* RNA FISH (n=5). Confocal images show *GZMB* RNA (A) or *IFNγ* RNA (B)-expressing CAR T cells in close proximity in a representative area close to the residual CD19+ tumor in a representative area close to the residual tumor. The whole tissue overview images of CAR RNA, CD19 with either *GZMB* RNA (C) or *IFNγ* RNA (D) staining shows the global distribution of the different markers. Whole tissue RRS analysis of the two RNA FISH, the CD19 IHC, the WGA and DAPI channel of each single tile was performed and the various x-y plots depict the spatial distribution of CD19+, CAR+ and *GZMB*+ (E) cells or *IFNγ*+ cells (F) within the tumor section. Scale bar = 10 μm

Around 2500 or 2800 CD19+ cells were detected in the *GZMB* and *IFNγ* datasets, respectively (Fig. 5A). We detected approximately >31000 CAR+ *GZMB-* and 700 CAR+ *GZMB+* cells/tissue section as well as >36000 CAR+*IFNγ*- and 40 CAR+ *IFNγ+* cells/tissue section (Fig. 5B). Therefore, the frequency and spatial density of CAR+ *GZMB+* cells was approximately 2% or 37 cells/mm^2^ whereas for CAR+ *IFNγ+* cells it was below 0.2% or 1.8 cells/mm^2^ (Fig. 5C, D). The frequency and spatial density of CAR+ cells that were negative for either *GZMB* or *IFNγ* was above 98% or around 1500 CAR+/mm^2^ (Fig. 5C, D).

**Figure 5:**
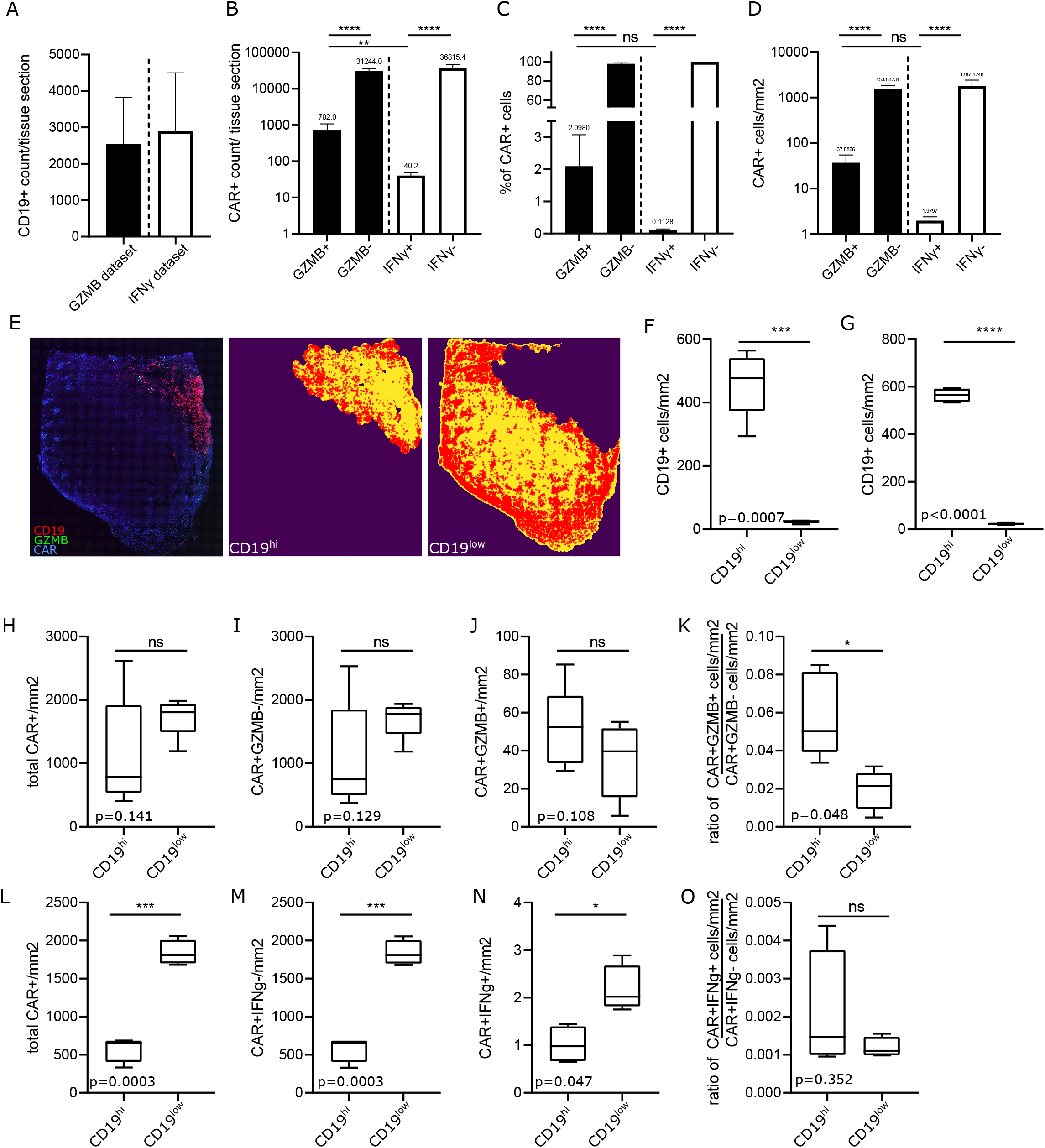
Spatial density analysis of the *GZMB+* and *IFNγ+* CAR T cells in CD19high and low areas reveals differential spatial effector gene expressio. Statistics of the CD19/CAR/*GZMB* (n=5) and CD19/CAR/*IFNγ* (n=5) cumulative datasets from Figure 4 and supplementary Fig. 8 and 9. Total CD19+ cells/mm2 (A) as well as total number (B), frequency (C) and spatial density (D) of CAR+ *GZMB*-, CAR+ *GZMB*+, CAR+ *IFNγ*-, CAR+ *IFNγ* + are shown. To assess the effect of CD19 antigen expression on CAR T cell function tumor sections were separated into a CD19high and CD19low areas and the number of CAR T cells that express *GZMB* and *IFNγ* or not within the two areas were counted. An example shows how tissue sections were separated in CD19high and CD19low areas denoted by the yellow color and, in this case the CAR+*GZMB*-cells as red dots (E). Spatial density of CD19+ cells in the CD19high and CD19low area in the CD19/CAR/*GZMB* (n=5) (F) and CD19/CAR/*IFNγ* (n=4) (G) cumulative datasets. For the CD19/CAR/*GZMB* cumulative dataset, total CAR+ cells/mm2 (H), CAR+ *GZMB*-cells/mm2 (I), CAR+ *GZMB*+ cells/mm2 (J), and the ratio of CAR+ *GZMB*+ to CAR+ *GZMB*-cells/mm2 (K) in the CD19high and CD19low area are shown. For the CD19/CAR/ *IFNγ* cumulative dataset, total CAR+ cells/mm2 (L), CAR+ *IFNγ-* cells/mm2 (M), CAR+ *IFNγ+* cells/mm2 (N), and the ratio of CAR+ *IFNγ+* to CAR+ *IFNγ-* cells/mm2 (O) in the CD19high and CD19low area are shown. C-D were analyzed by one-way ANOVA. F-O were analyzed by paired t test.

To analyze the effect of tumor antigen expression on the spatial expression of effector gene *GZMB* and *IFNγ,* the spatial datasets were divided in a CD19high and CD19low area (Fig. 5E). The spatial density of CD19+ cells/mm^2^ in the CD19high and CD19low area in the *GZMB* and *IFNγ* dataset were significantly different (Fig. 5F, G) indicating that CAR T cells had a higher probability for antigen exposure in the CD19high area. For the *GZMB* dataset, the number of total CAR+ (Fig. 5H), CAR+ *GZMB-* (Fig. 5I) and CAR+ *GZMB+* cells/mm^2^ (Fig. 5J) was not significantly different between the CD19high and CD19low area, however, there was a trend towards higher spatial density of total CAR+ and CAR+ *GZMB*-cells/mm^2^ in the CD19low area and an opposite trend of higher spatial density for CAR+*GZMB*+ cells/mm^2^ in the CD19high area. The ratio of CAR+ *GZMB*+ to CAR+ *GZMB*-cells/mm^2^ was significantly higher in the CD19high area compared to the CD19low area indicating an association of GZMB expression and the presence of the cognate CAR antigen (Fig. 5K). For the *IFNγ* dataset, the number of total CAR+ (Fig. 5L), CAR+ *IFNγ-* (Fig. 5M) and CAR+ *IFNγ+* cells/mm^2^ (Fig. 5N) was significantly higher in the CD19low area compared to the CD19high area. The ratio of CAR+ *IFNγ*+ to CAR+ *IFNγ*-cells/mm^2^ was not significantly different in a comparison of the CD19high and CD19low area (Fig. 5O). In summary, these data indicate that the RRS CAR RNA FISH approach in combination with RNA FISH for T cell lineage and effector genes can provide important insights into the spatial distribution, functionality and heterogeneity of CAR T cells on a single cell level.

## Discussion

In this study, we present the development of a multiplex RNA FISH IHC assay that is coupled with a RRS algorithm for image segmentation and quantitation of engineered cells. This assay enables to interrogate single cell expression levels of multiple mRNA of interest such as *CD4*, *Cd8α*, *IFNγ* and *GZMB* and their spatial distribution and can be used to detect, quantitate and evaluate the frequency and function of CAR T cells and cancer cells in tissues. The use of specific probes directed against both the wPRE and scFv allows accurate detection of CAR in tissues. It circumvents the common caveats of immunohistochemistry such as long lead-time for antibody development and overlap of endogenous epitopes with the chimeric protein of interest. This approach can be quickly modified for *in situ* trafficking studies for novel CARs with different target proteins, e.g. CD20, CD22, BMCA or any other genetically modified cells *(18)* and does not require the addition of fluorescent protein or tracking molecules to the CAR expression cassette. It can be used to study the interplay of CAR T cells, tumor cells and the tumor microenvironment in solid tumor to develop novel strategies and hypothesis to improve CAR T cell function in solid tumors; an area where CAR T cells therapies still fall short of the clinical responses seen in hematologic cancers *(18, 19)*.

To know the spatial distribution and phenotype of CAR T cells is likely relevant to optimize this promising cellular therapy for solid tumors. This is exemplified by the functional and spatial heterogeneity of CAR T cells that express genes associated with cytotoxicity such *GZMB* or *IFNγ* with respect to the residual solid tumor and adjacent tissue. The decreased *GZMB* expression in distant, CD19 negative tissues suggests cell-to-cell contact with target cells is required for CAR T-cell activation. In contrast to the *GZMB+* CAR T cells, the more delocalized expression of *IFNγ* may indicate that these CAR T cells migrate in and out from the area of the tumor surface more frequently or express *IFNγ* for an extended period of time post-antigen exposure. Furthermore, there is still a considerable number of CAR T cells close to the residual tumor surface that does not express *IFNγ* or *GZMB.* Further studies into CAR T cell functional heterogeneity in xenograft models and human tumor and healthy tissue and how it relates to clinical response rates will likely contribute to improvement of future CAR T cells interventions.

Globally, this assay can provide insight on how CAR T cells are acting within a solid tumor setting and how the heterogeneity effector genes such as *GZMB* and *IFNγ* of the CAR T cell population affects the reduction in tumor size. The IFNγ-expressing CAR T cells may change the local tumor microenvironment such as M2-to-M1 macrophage differentiation or stimulate incoming tumor-infiltrating T cells and, thus gradually lead to tumor regression. However, extended IFNγ-expression from tumor-infiltrating T lymphocytes is associated with increase PD-L1 expression and immune evasion in melanoma *(20)* and neuroblastoma *(21)* and thus, extended IFNγ secretion from CAR T cells may be also detrimental in a solid tumor setting. Of note, high levels of serum IFNγ are also implicated in an increased risk of grade 3–5 neurotoxicity in anti-CD19 CAR T cell therapy *(5, 22)*. Further evaluation of *in situ* cytokine-chemokine expression of CAR T cells in tissue is likely to lead to increased understanding of the mechanism of CAR T cell tumor killing and toxicity.

Quantitative PCR, flow cytometry and bioluminescence imaging are the main tools to study CAR T cells in vivo pre-clinical and, to some extent, clinical studies. Even though these assays allow high sample throughput, macroscopic analysis of the tumor mass and/or provide high statistical power for quantification, none of them assesses the spatial distribution CAR T cells and the interplay of CAR T cells and their targets.

The RRS CAR RNA FISH approach described here provides an easy-to-go tool kit for the CAR T cells studies and promises to provide insights into the trafficking, persistence and phenotype of CAR T cells in liquid and solid tumors in preclinical models to better understand the interplay between the tumor microenvironment and CAR T cell function.

## Acknowledgements

We thank S. Sun, L. Schroeder, P Guo, J. Vazquez and D. McDonald for technical support and Mindy Miner for editing the manuscript.

This independent research was supported by National Institute of Health grant UM1 A126623 and by the Gilead Sciences HIV Cure Grant Program.

This research was supported by the Experimental Histopathology Shared Resource of the Fred Hutch/University of Washington Cancer Consortium (P30 CA015704).

This research was supported by the Cellular Imaging Shared Resource (CISR) of the Fred Hutch/University of Washington Cancer Consortium (P30 CA015704).

The authors declare no competing interest.

## Code and data availability statement

Code and raw data are available upon request

## Author contributions

K.E. and L.C. conceived the project. K.E. performed most experiments, analyzed the data together with A.L. and interpreted the data together with LC.

A.L. developed the algorithm and A. L. and K.E. performed the RRS analysis.

K.D. helped with experiments.

J.Z. helped with scientific advice and valuable expertise

M.J. provided tissue samples and scientific advice.

KE and LC wrote the manuscript with input from all authors.

## Figures

**Supplementary Figure 1: CAR probe table, principle of RNA FISH.**

Three zz probes sets were designed for detection of anti-CD19 CAR T cells. The probes bind to the scFv, the intergeneic regions between the signaling domains and some probes sets to the 3′ untranslated region of the CAR mRNA, which constitute nucleic acid sequences that cannot be found in non-genetically engineered organisms (A).

**Supplementary Figure 2: purity of primary CAR T cell culture after expansion.**

Anti-CD19 CD8 CAR T cells were cultured with γ-irradiated CD19+ TM-LCL cells to stimulate expansion. On day 12 of cell culture, CAR T cell purity was assessed by flow cytometry for the cell surface markers CD19-BV786, EGFRt-PE and CD8-FITC (n=1). These cells were processed for CAR RNA FISH and used for the *in vitro* validation.

**Supplementary Figure 3: CAR RNA FISH detects CAR RNA mainly extranuclear.**

CAR Jurkat cells were stained with CARprobes and nuclear counterstain to exemplify subcellular localization of CAR RNA. The CAR RNA is mainly detected extranuclear or on the outline of the nuclear counterstain.

**Supplementary Figure 4: presence and expression of CAR sequence and wPRE in mouse xenografts.**

ddPCR analysis for the presence of the CAR signaling domain (CD8CD3) and the 3′UTR containing the wPRE sequence of genomic DNA in xenograft tissue section from CAR T cell-infused and control mice. (n=2). ddPCR primers and probes were directed against the CD3 CD8 region and wPRE portion to test for CAR and CD19 lentivirus-derived expression cassettes.

**Supplementary Figure 5: CAR RNA FISH in situ validation.**

Specificity of CAR RNA FISH in situ staining was further validated with RNA FISH staining for housekeeping gene *PPIB* (A) and bacterial gene *DapB* (B) CAR RNA FISH protocol without the addition of CAR FMZ63 15zz probes (C) or after RNAse treatment (D) (n=3).

**Supplementary Figure 6: Comparison of manual and RRS cell counting**

We chose 10 images from the CAR+ BE2 tumor dataset that are exemplary for images without artifacts or that show different imaging artifacts like strong membrane staining or weak nuclei staining (A). Total cell numbers were counted manually by three operators or with the RRS algorithm using the nuclei and membrane WGA staining (B). Correlation of the number of cells counted/images by the three operators and the RRS method (C). The counting time required/image is shown in (D).

**Supplementary Figure 7: Low effector gene expression in CAR T cells that are located further in the CD19low area.**

CAR T cell functional phenotyping in a BE2 neuroblastoma xenograft was done by IHC for tumor marker CD19 and CAR RNA FISH in combination with either *GZMB* (n=5) or *IFNγ*RNA FISH (n=5). Confocal images of the combination of CAR RNA and CD19 with either *GZMB* RNA (A) or *IFNγ* RNA (B) in a representative area in the CD19low area. Exemplary wide field images that were used to create the overview thumbnail images in Figure 4 image of CAR RNA, CD19 with either *GZMB* RNA (C) or *IFNγ* RNA (D).

**Supplementary Figure 8: RRS analysis CD19, CAR and *GZMB* expression of tissue sections from a BE2 tumor.**

CAR T cell functional phenotyping on consecutive section of a BE2 neuroblastoma xenograft was done by counterstaining for nuclei and membranes (not shown) as well as IHC for tumor marker CD19 and CAR RNA FISH in combination with *GZMB* RNA FISH. The images were analyzed by RRS and plotted in FlowJo(n=).

**Supplementary Figure 9: RRS analysis CD19, CAR and *IFNγ* expression of tissue sections from a BE2 tumor.**

CAR T cell functional phenotyping on consecutive section of a BE2 neuroblastoma xenograft was done by counterstaining for nuclei and membranes (not shown) as well as IHC for tumor marker CD19 and CAR RNA FISH in combination with *IFNγ* RNA FISH. The images were analyzed by RRS and plotted in FlowJo(n=4).

**Supplementary Table 1:** CAR RNA FISH probes and respective catalog number.

| Probe name | Catalog Number |
| --- | --- |
| FMC63 15ZZ | 515321 |
| FMC63 26ZZ | 513551 |
| DapB-C1 | 310043 |
| Hs-PPIB-C2 | 313901-C2 |
| Hs-PPIB-C1 | 313901-C1 |
| Hs-GZMB | 445971 |
| Hs-GZMB-C2 | 445971-C2 |
| Hs-CD8A-C3 | 560391-C3 |
| Hs-CD4-C2 | 605601-C2 |
| Hs- $IFN\gamma$-C1 | 310501-C1 |
| Hs- $IFN\gamma$-C2 | 310501-C2 |

