## Supplementary Figures for "A CAR RNA FISH assay to study functional and spatial heterogeneity of chimeric antigen receptor T cells in tissue"

| | total zz | scFv | CAR core | 3'UTR | target tissue/species |
|---|---|---|---|---|---|
| FMC63 15zz | 15 | 11 | 4 | 0 | human and mouse tissue containing different lentiviral constructs |
| FMC63 27zz | 26 | 11 | 4 | 12 | human and mouse tissue with only CAR T cells present |

Supplementary Figure 1

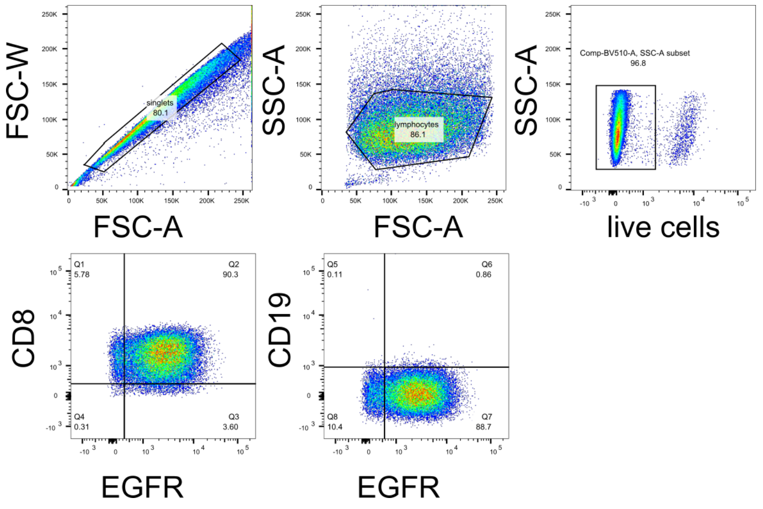


Supplementary Figure 2

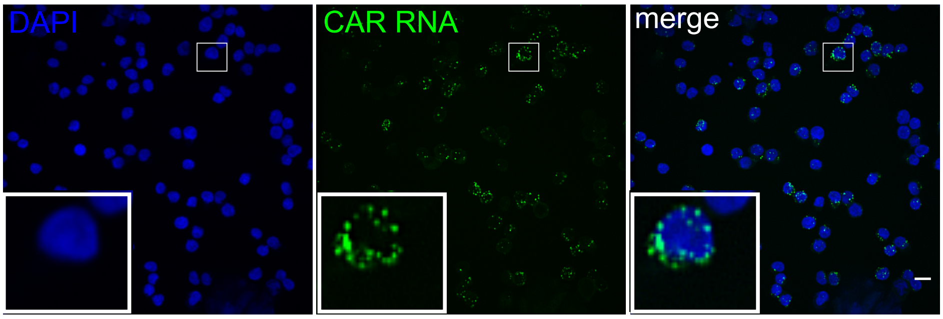


Supplementary Figure 3

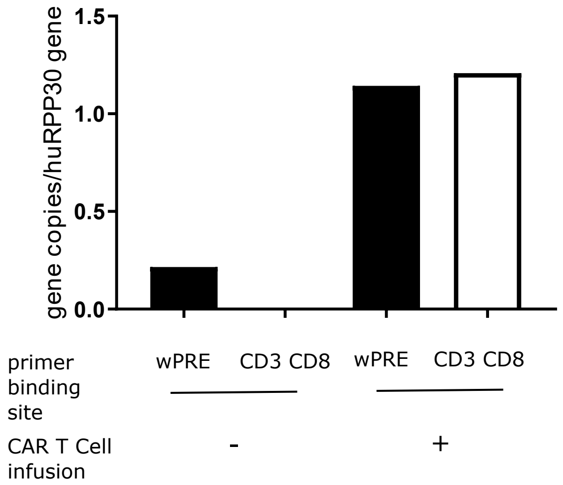


Supplementary Figure 4

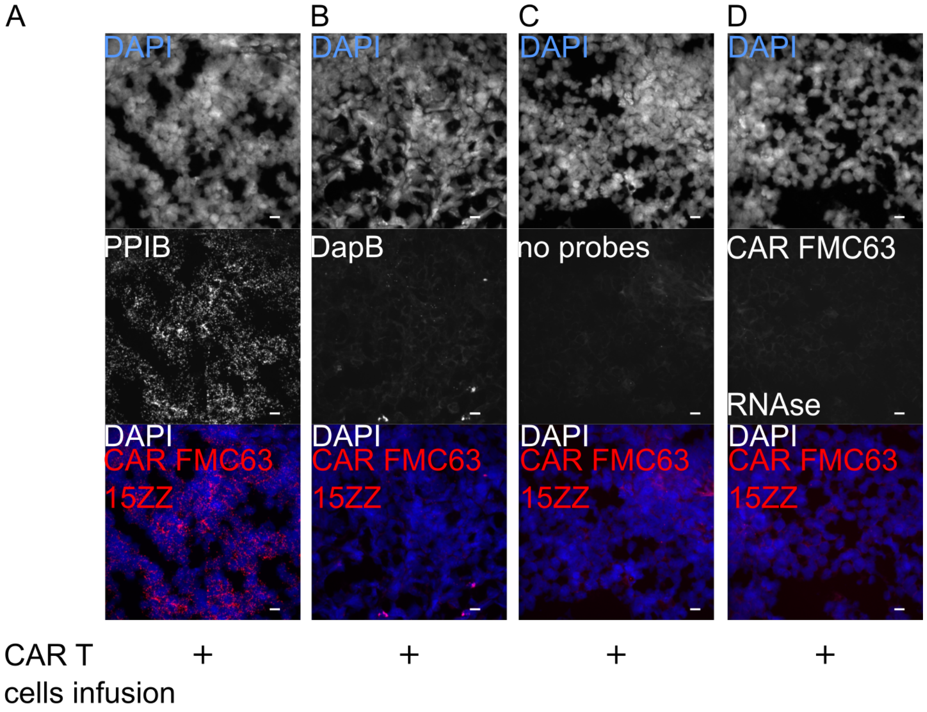


Supplementary Figure 5

A

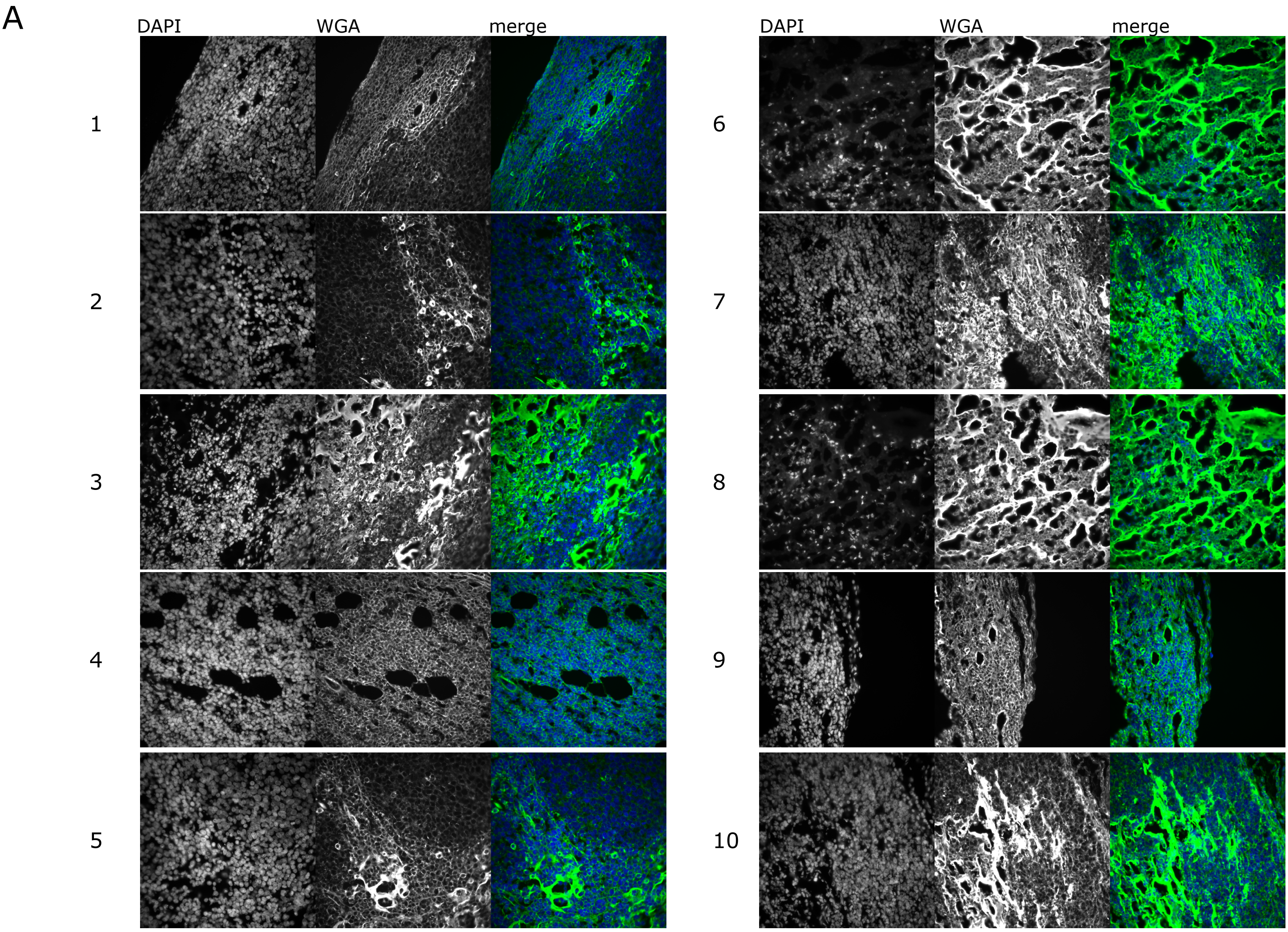


B

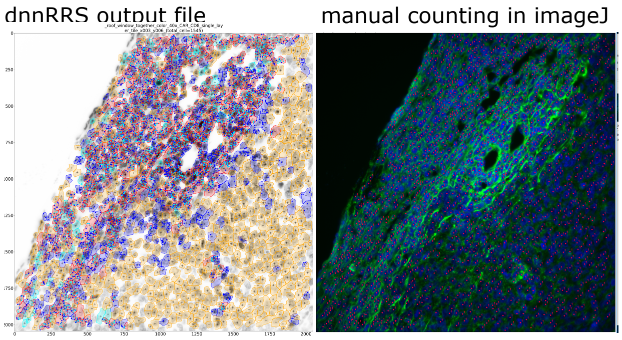


C

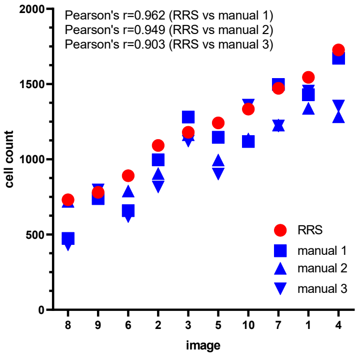


D

| image | dnnRRS (h:mm:ss) | manual 1 (h:mm:ss) | manual 2 (h:mm:ss) | manual 3 (h:mm:ss) |
|---|---|---|---|---|
| 1 | 0:01:41 | 0:14:50 | 0:12:26 | 0:41:19 |
| 2 | 0:01:41 | 0:08:49 | 0:07:29 | 0:28:44 |
| 3 | 0:01:43 | 0:11:27 | 0:10:06 | 0:29:55 |
| 4 | 0:01:27 | 0:14:56 | 0:11:25 | 0:23:34 |
| 5 | 0:01:49 | 0:10:16 | 0:09:16 | 0:13:40 |
| 6 | 0:01:32 | 0:15:01 | 0:07:57 | 0:11:37 |
| 7 | 0:01:38 | 0:14:02 | 0:10:47 | 0:18:37 |
| 8 | 0:00:58 | 0:10:56 | 0:07:09 | 0:25:23 |
| 9 | 0:01:07 | 0:12:35 | 0:08:09 | 0:15:41 |
| 10 | 0:01:06 | 0:20:46 | 0:12:45 | 0:26:03 |
| avergage time/image | 0:01:28 | | 0:15:31 | |
| SD | 0:00:18 | | 0:08:02 | |

Supplementary Figure 6

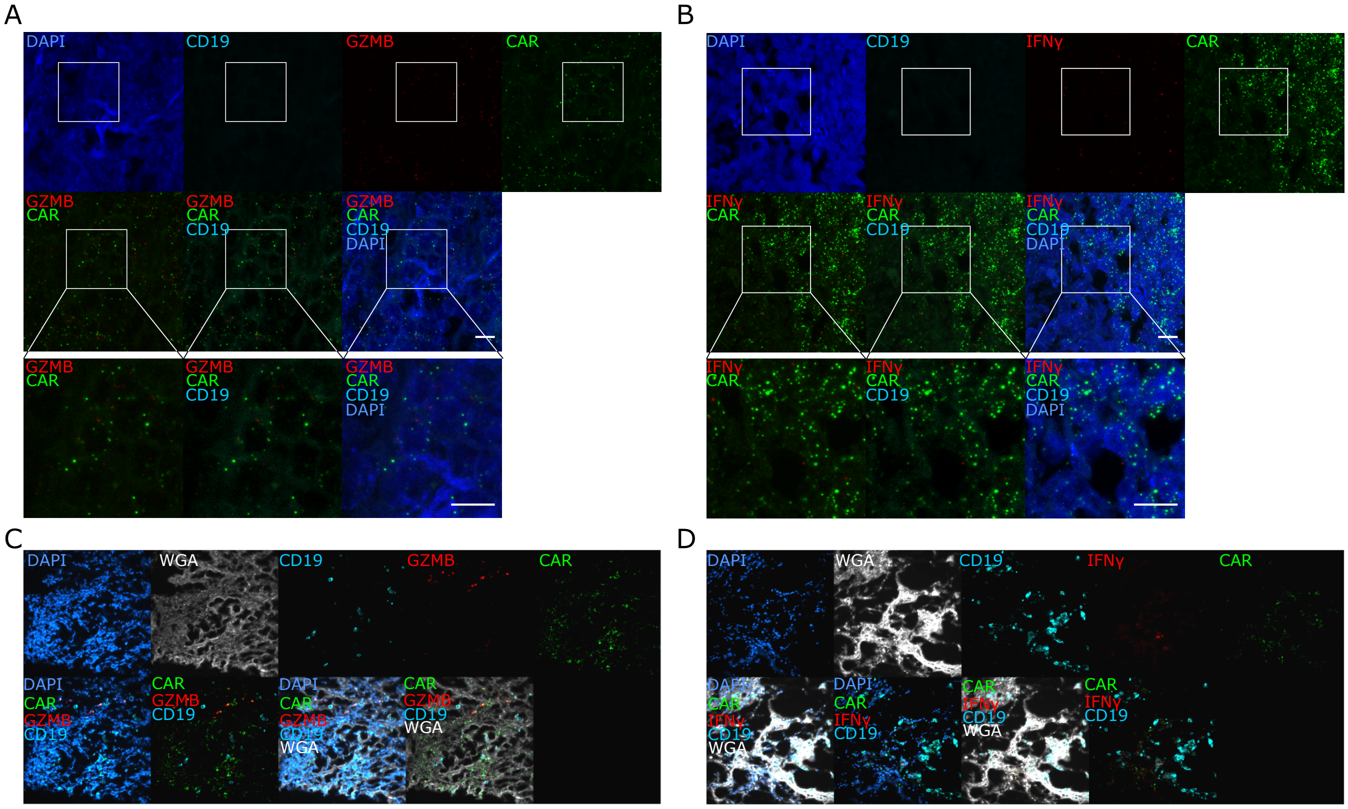


Supplementary Figure 7

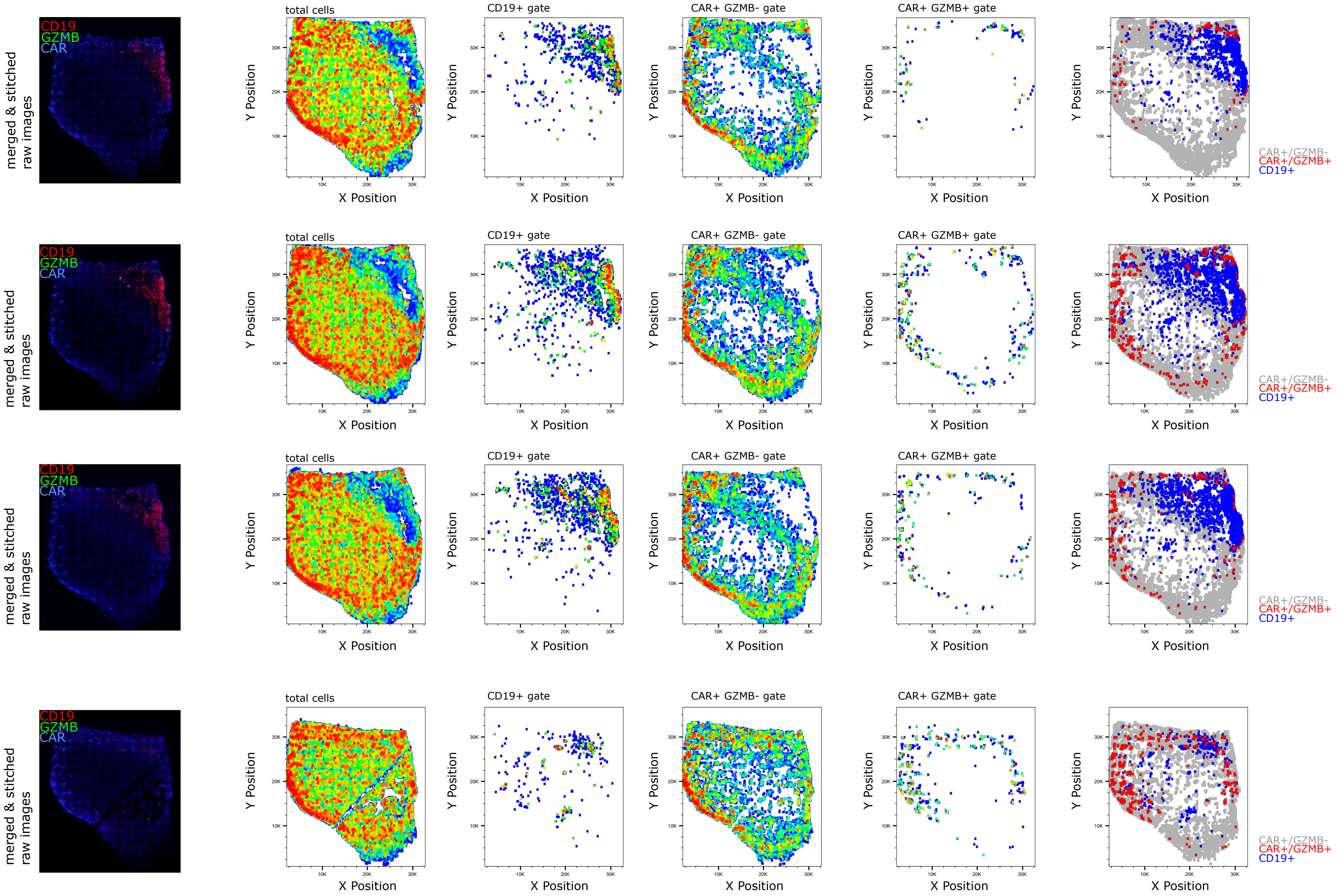


Supplementary Figure 8

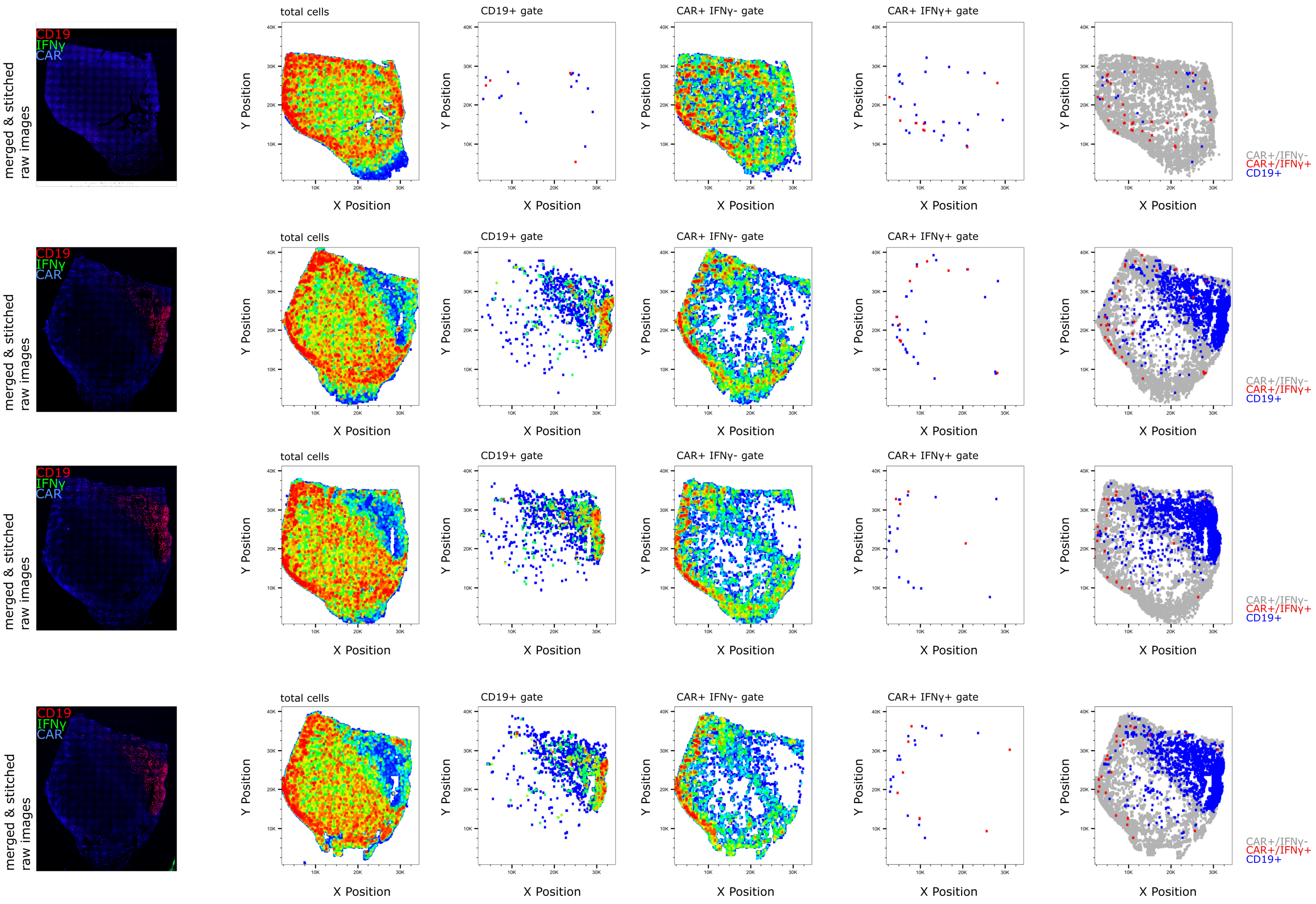


Supplementary Figure 9
